# OPTICAL BLUR AFFECTS DIFFERENTLY ON AND OFF VISUAL PATHWAYS

**DOI:** 10.1101/2024.10.17.618707

**Authors:** Carmen Pons, Reece Mazade, Jianzhong Jin, Mitchell Dul, Jose-Manuel Alonso

**Author notes:** Send correspondence to: Jose-Manuel Alonso SUNY Optometry Department of Visual Sciences 33 West, 42nd street, 17th floor New York, NY 10036 Voice.

## Abstract

The human eye has a crystalline lens that focuses retinal images at the point of fixation. Outside this fixation region, images are distorted by optical blur, which increases light scatter and reduces the spatial resolution and contrast processed by neuronal pathways. The spectacle lenses that humans use for optical correction also minify or magnify the images, affecting neuronal surround suppression in visual processing. Because light and dark stimuli are processed with ON and OFF pathways that have different spatial resolution, contrast sensitivity and surround suppression, optical blur and image magnification should affect differently the two pathways and the perception of lights and darks. Our results provide support for this prediction in cats and humans. We demonstrate that optical blur expands ON receptive fields while shrinking OFF receptive fields, as expected from the expansion of light stimuli and shrinkage of dark stimuli with light scatter. Spectacle-induced image magnification also shrinks OFF more than ON receptive fields, as expected from the stronger surround suppression in OFF than ON pathways. Optical blur also decreases the population response of OFF more than ON pathways, consistent with the different effects of light scatter on dark and light stimuli and the ON-OFF pathway differences in contrast sensitivity. Based on these results, we conclude that optical blur and image magnification reduce the receptive field sizes and cortical responses of OFF more than ON pathways, making the ON-OFF response balance a reliable signal to optimize the size and quality of the retinal image.

**HIGHLIGHTS:**

1. Optical blur affects ON and OFF pathways differently.
2. Blur expands ON but shrinks OFF receptive fields, weakening OFF more than ON signals.
3. Magnification from positive blur reduces OFF more than ON signal strength and latency.
4. ON-OFF response balance can signal retinal image quality and guide eye growth.

**IN BRIEF:** Pons et al. demonstrate that optical blur affects differently the receptive field properties and responses from ON and OFF visual pathways, revealing a new relation between ON/OFF pathway balance and blurred visual perception. The results highlight a new role of ON and OFF pathways in optimizing retinal image quality that can have implications for the regulation of eye growth.

**GRAPHICAL ABSTRACT:** 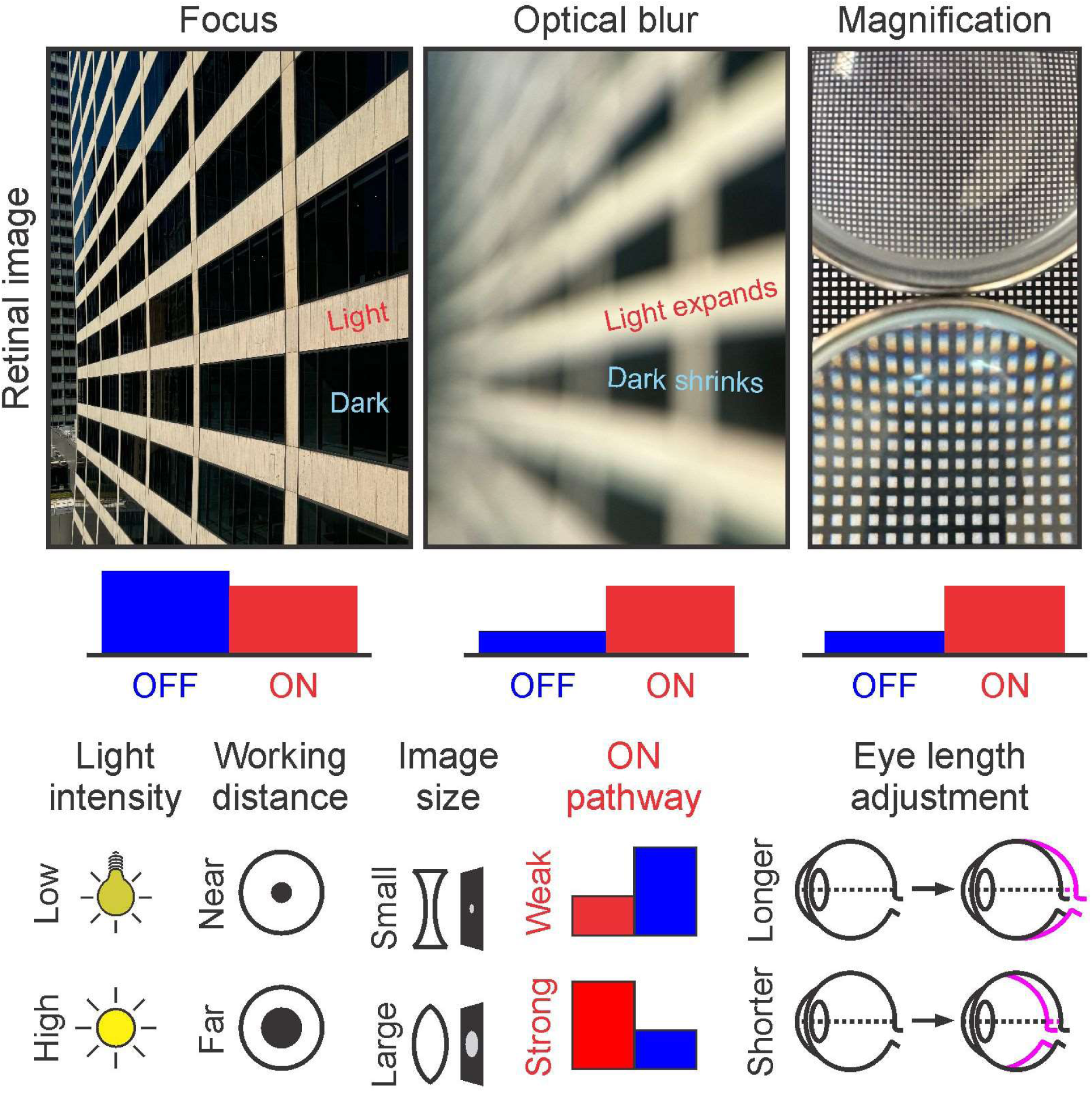

## INTRODUCTION

Optical blur plays an important role in the development of the visual pathway. In humans, a mismatch between the optical power of the eye and its axial length can lead to nearsightedness (blurred vision at far distance) or farsightedness (blurred vision at near distance). These two visual disorders, also known as myopia and hyperopia, can be induced in a large variety of animal models by exposing the eye to negative or positive optical blur ^1,2^. Exposure to optical blur also affects the neuronal wiring of the brain ^3–7^ and can disrupt the processing of light and dark stimuli mediated by ON and OFF visual pathways ^8–10^.

Optical blur causes light scatter, which distorts the size, contrast and spatial resolution of stimuli. Because ON and OFF pathways have different surround suppression, contrast sensitivity, and spatial resolution ^11–25^, they should be differently affected by optical blur. The increase in light scatter should expand the size of light stimuli while shrinking the size of dark stimuli, causing opposite distortions in the size tuning of ON and OFF pathways. In addition, the contrast reduction should affect OFF more than ON pathways because ON-pathway responses saturate more with contrast and are therefore more resistant to contrast loss ^8,10,12,18,26,27^. The image magnification caused by positive spectacles should also affect OFF more than ON pathways because OFF pathways have stronger surround suppression ^11,19,20^. Only the contrast loss of the finest image details should affect ON more than OFF pathways because ON pathways respond to higher spatial frequencies than OFF pathways ^13,18,28^.

Our cortical measurements confirm these predictions while revealing a new relation between ON-OFF pathway asymmetries and blurred visual perception ^8,9^. They also provide new insights on the role of ON and OFF pathways in signaling image size and sharpness during eye development.

## RESULTS

We performed electrophysiological recordings from the visual cortex of cats and humans wearing contact lenses or spectacles of different optical power while having their accommodation pharmacologically blocked. In cat primary visual cortex, we performed multielectrode recordings of multiunit activity to measure the effect of optical blur on cortical receptive field size and visual responses to diverse stimuli. In human visual cortex, we performed electroencephalography recordings to measure the effect of optical blur on responses to light and dark checkerboard stimuli. Our results demonstrate that optical blur affects the visual responses of ON and OFF cortical pathways differently in both cats and humans.

### Optical blur expands ON receptive fields while shrinking OFF receptive fields

We investigated how optical blur affects ON and OFF cortical receptive fields in cat visual cortex. Optical blur induced with contact lenses made ON receptive fields larger and OFF receptive fields smaller, as expected from the effect of light scatter, which expands light stimuli and shrinks dark stimuli (Figure 1a-b). This expansion of ON receptive fields and shrinkage of OFF receptive fields could be demonstrated with both negative and positive blur. In contrast, spectacles placed ∼1 cm in front of the eye shrunk OFF receptive fields only with positive blur (Figure 1c).

**Figure 1.**
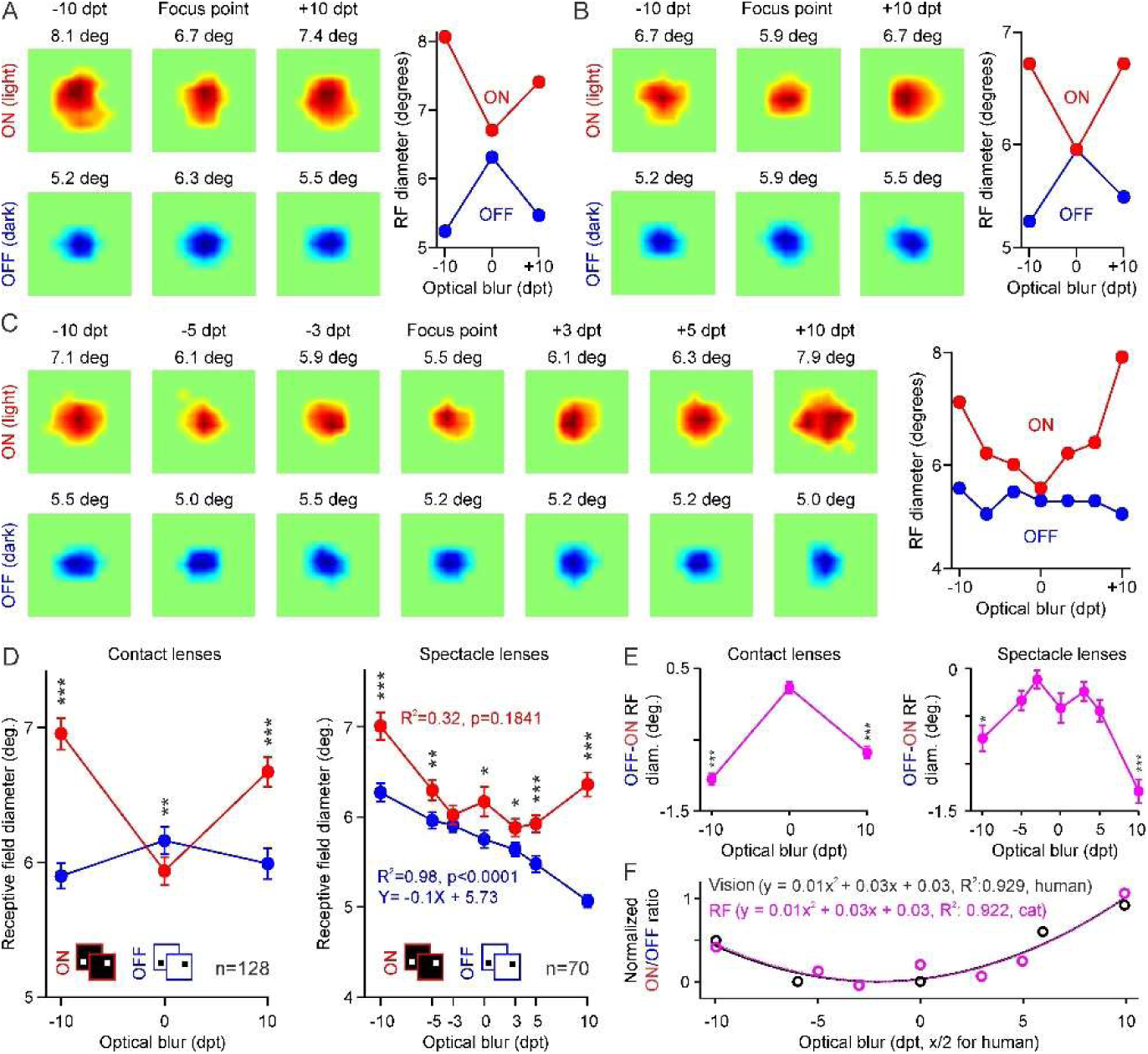
Optical blur and image magnification affect differently the ON and OFF visual pathways. **(A)** Left: Receptive fields mapped with light (top, red) and dark stimuli (bottom, blue) while inducing optical blur with contact lenses. Top labels: optical blur of -10, 0, or +10 diopters and receptive field diameter (deg: degrees). Right: Change in receptive field diameter with optical blur. (**B)** Same as in A but for a different cortical recording site. (**C)** Same as A-B but inducing optical blur with spectacle lenses (-10, -5, -3, 0, +3, +5, +10 diopters). **(D)** Left: Average receptive field diameter (n=128, 88 OFF-dominated) measured with dark (top, blue) and light stimuli (top, red) while inducing optical blur with contact lenses. The text label reports goodness of fit (R^2^) and p values of linear regressions (equations provided only for significant correlations). Right: Same as in left panel, but inducing optical blur with spectacles (n=70, 43 OFF-dominated). **(E)** Same as D for OFF-ON difference in receptive field (RF) diameter (diam.). **(F)** Same data as E right panel, but plotting normalized ON/OFF RF diameter ratio (magenta) superimposed with light/dark ratio of perceptual errors measured in humans (black, data from Pons et al. (2017)). Both measurements were fit with quadratic equations, normalized (subtracting minimum and dividing by maximum of the fit), and shown superimposed after multiplying human optical blur by 2 (cats have lower spatial resolution than humans and their receptive field eccentricity is 5-10 degrees, not zero). Labels at the top report the equations and R^2^. * p<0.05, ** p<0.01, *** p<0.001 (Wilcoxon tests). Error bars indicate ± SEM.

On average across multiple cortical neurons and blur levels, ON receptive fields were significantly larger than OFF receptive fields (Figure 1d, mean ± SEM measured with contact lenses: 6.52°±0.07° for ON vs. 6.02°±0.06° for OFF, n=128, p<0.00001; spectacles: 6.24°±0.05° for ON vs. 5.73°±0.04° for OFF, n= 70, p<0.00001, Wilcoxon sign-rank tests), and the ON-OFF size difference increased with the absolute magnitude of optical blur (Figure 1e, ON-OFF: -0.23°± 0.08° at 0 diopters vs. 0.87°±0.09° at ±10 diopters, p<0.00001 for contact lenses; 0.41°±0.16° at 0 diopters vs. 1.02°±0.13° at ±10 diopters, p=0.00009 for spectacles, Wilcoxon sign-rank tests). At focus, the average OFF receptive field could be slightly larger (Figure 1d, left) or smaller than the average ON receptive field (Figure 1d, right), depending on the number/strength of the OFF cortical domains included in the sample (receptive fields in OFF cortical domains have larger OFF than ON subregions ^15,16,29–31^). Furthermore, the effect of optical blur on cortical receptive field size measured in cats shared a striking resemblance (Figure 2f) with the effect of optical blur measured in human visual perception (data from^8^).

**Figure 2.**
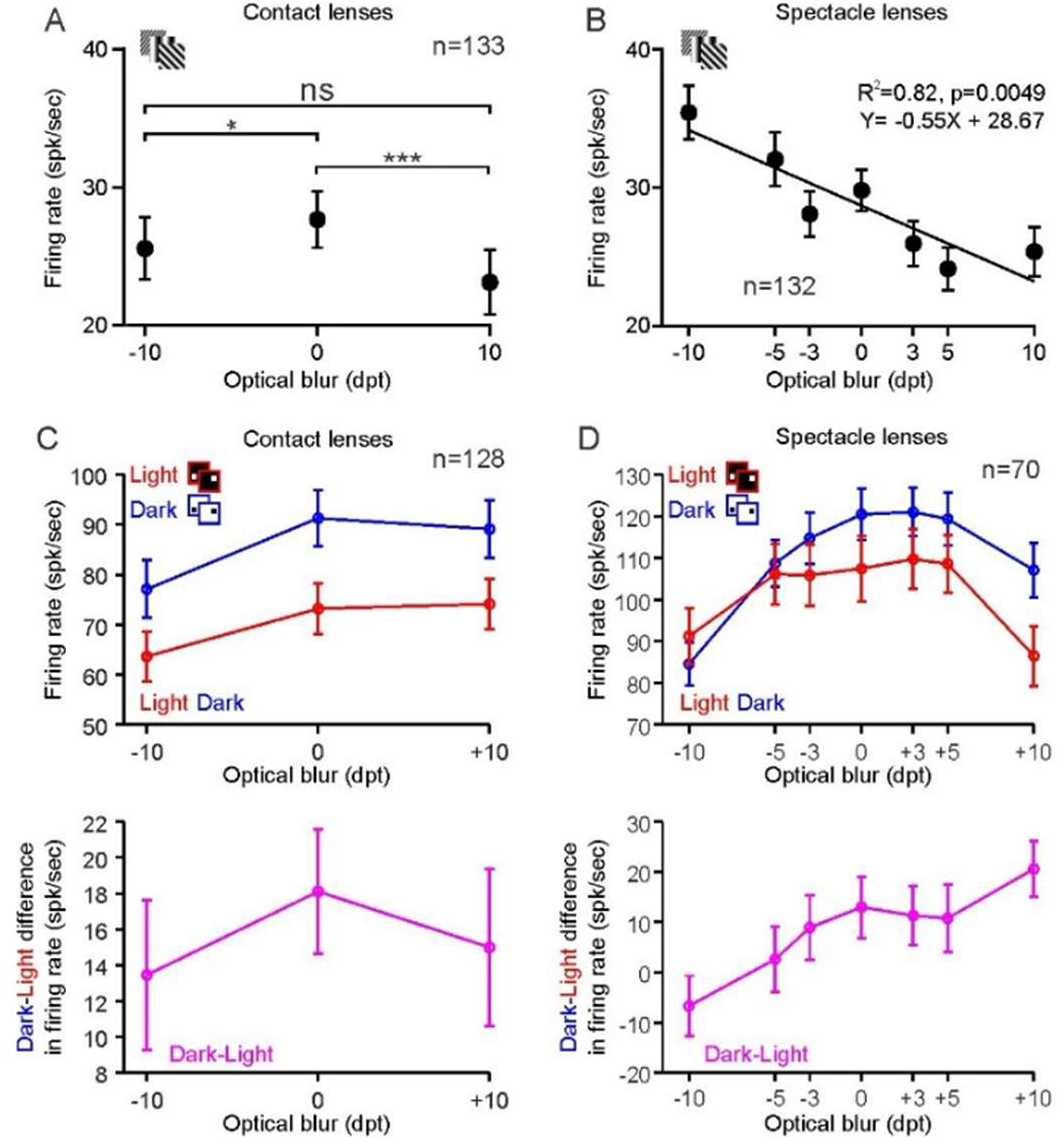
Positive blur and spectacle image magnification reduce cortical responses to grating patterns and increase OFF pathway dominance to small stimuli. **(A)** Average response to large grating patterns measured while inducing optical blur with contact lenses (n=133). **(B)** Same as A for spectacle lenses (n=132). The line shows a linear regression fit to the data (text label reports equation, R^2^, and statistical significance). **(C)** Top: same as A for responses to small dark (blue) and light (red) stimuli measured while inducing optical blur with contact lenses (n=128). Bottom: same as top for dark-light difference (magenta). **(D)** Same as C but for spectacle lenses (n=70). * p<0.05, ** p<0.01, *** p<0.001 (Wilcoxon test). Error bars indicate ± SEM.

### Spectacle image magnification reduces cortical receptive field size

When induced with contact lenses, negative blur expanded ON receptive fields slightly more than positive blur (Figure 1d, left, -10 vs. +10 diopters: 6.95°±0.12° vs. 6.67°±0.11°, p=0.00002; ±10 vs. 0 diopters: 6.81°±1.23° vs. 5.94°±1.18°, p=2.13x10^-18^, Sign-rank Wilcoxon test), but both blur polarities shrunk similarly OFF receptive fields (Figure 1d, left, -10 vs. +10 diopters: 5.90°±0.09° vs. 5.99°±0.09°, p=0.0914; ±10 vs. 0 diopters: 5.95°±1.14° vs. 6.16°±1.16°, p=1.2x10^-5^, Sign-rank Wilcoxon tests). By comparison, when induced with spectacles, negative blur expanded ON receptive fields more than positive blur (Figure 1d, right, -10 vs. +10 diopters: 7.01°±1.28° vs. 6.36°±1.11°, p=0.0016; ±10 vs. 0 diopters: 6.69°±0.98° vs. 6.17°±1.34°, p=9.15x10^-5^, Sign-rank Wilcoxon test), but OFF receptive fields only shrunk with positive blur (Figure 1d, left, -10 vs. 0 diopters: 6.28°±0.86° vs. 5.76°±0.80°, p=1.04x10^-9^; +10 vs. 0 diopters: 5.07°±0.59° vs. 5.76°±0.80°, p=1.35x10^-10^, Sign-rank Wilcoxon test).

The image magnification of positive spectacles also made OFF receptive field size strongly correlated with spectacle blur (R^2^=0.98; p<0.0001). As spectacle blur became more positive and image magnification increased, dark stimuli became larger and more effective at driving surround suppression, making OFF receptive fields smaller (the receptive fields also became smaller because of the sampling decrease with larger stimuli). Consistently with the stronger surround suppression in OFF than ON pathways ^11,19,20^, the effect of image magnification was also stronger for dark than light stimuli (Figure 1d, right, -10/+10 ratio for OFF vs. ON: 1.24±0.02 vs. 1.12±0.03, p=5.95x10^-5^, Sign-rank Wilcoxon test). It is important to notice that, whereas spectacles with positive blur caused a reliable and systematic decrease in OFF receptive field size (R^2^ = 0.98), the reduction was extremely small. On average, the OFF receptive field size decreased by just 0.1 degree per diopter, which is 1/60 (< 2%) of the average receptive field size measured in our sample (∼6°). Therefore, we conclude that positive spectacle blur causes a very small but reliable decrease in cortical receptive field size because it magnifies stimuli. This effect may explain why human subjects wearing positive (or negative) spectacles perceive a noticeable increase (or decrease) in image size.

### Optical blur and image magnification weaken cortical responses

Optical blur also reduced the strength of cortical responses to large grating patterns. When optical blur was induced with contact lenses, the responses to gratings decreased by ∼12% (Figure 2a, 0 vs. ±10 diopters: 27.65±2.05 vs. 24.33±1.62 spk/sec, p=0.0012, Rank-sum Wilcoxon test) and the reduction was not significantly different between negative and positive blur (Figure 2a, -10 vs. +10 diopters: 25.55±2.25 vs. 23.11±2.34 spk/sec, p = 0.9508, Sign-rank Wilcoxon test). Conversely, when optical blur was induced with spectacle lenses, the responses to gratings decreased with positive blur but increased with negative blur, as expected from the effect of image magnification/minification on surround suppression (Figure 2b, 0 vs. +10 diopters: 29.78±1.50 vs. 25.35±1.78 spk/sec, p=0.005; 0 vs. -10 diopters: 29.78±1.50 vs. 35.40±1.92 spk/sec, p<0.00001, Sign-rank Wilcoxon tests). These magnification/minification effects made spectacle blur correlated with cortical response strength (Figure 2b, R^2^ =0.82, p=0.0049).

Optical blur also reduced cortical responses to small stimuli (Figure 2c, d). The response reduction was ∼6-9% with contact lenses (Figure 2c, 0 vs. ±10 diopters: 73.17±5.08 vs. 68.86±4.93 spk/sec for ON, p=0.0126; 91.27±5.55 vs. 83.08±5.56 spk/sec for OFF, p<0.00001, Wilcoxon tests) and ∼17-21% with spectacles (Figure 2d, 107.44±7.81 vs. 88.81±6.51 spk/sec for ON, p<0.00001; 120.43±6.20 vs. 95.76±5.10 spk/sec for OFF, p<0.00001, Wilcoxon tests). Optical blur induced with spectacles was strongly correlated with the response difference between dark and light stimuli (Figure 2d, bottom, R^2^=0.86, p=0.0025), a correlation that can be explained by the combined effect of image magnification and light scatter. That is, as spectacle blur becomes less negative (e.g. from -10 to -5 diopters), the reduction in image minification increases the stimulus size and spatial summation within the receptive field center making the responses stronger (Figure 2d, top). However, because the reduction in light scatter expands dark stimuli while shrinking light stimuli, the facilitation is more pronounced for dark stimuli and the dark-light response difference increases (Figure 2d, bottom). Similarly, as spectacle blur becomes more positive (e.g. from +5 to +10 diopters), the increase in image magnification enlarges the stimuli within the suppressive surround and reduces the response strength (Figure 2d, top). However, because the increase in light scatter shrinks dark stimuli while expanding light stimuli, the suppressive effect is more pronounced for light than dark stimuli, and the dark-light response difference still increases. Across blur levels, responses were also stronger to small dark than light stimuli as expected from the stronger responses from OFF pathways in visual cortex ^13–15,17,18,24,29,30,32–34^.

### Positive spectacle lenses decrease cortical response latency

Positive spectacle lenses also had a significant effect on cortical response latency (Figure 3). The response latency was shorter in OFF than ON cortical pathways (see also ^16,19,35^), and the difference could be demonstrated with both preferred and non-preferred stimuli (Figure 3a-b, OFF vs. ON: 44.25±0.66 ms vs. 46.55±0.60 ms for response enhancement, p=0.0045; 44.46±1.01 vs. 48.64±0.85 for response suppression, p=0.0011; Rank-sum Wilcoxon tests). The response latency to preferred stimuli was significantly correlated with spectacle blur in OFF but not ON cortical pathways (Figure 3c, R^2^=0.93, p=0.0004 for OFF versus R^2^ =0.25, p=0.2576 for ON), whereas the suppression latency to non-preferred stimuli was significantly correlated with spectacle blur in both pathways (Figure 3d, R^2^=0.65, p=0.0286 for OFF; R^2^=0.84, p=0.0038 for ON). Because increasing stimulus size reduces response latency in visual cortex ^19,20^, the decrease in response latency with spectacle blur can be explained by the increase in image magnification. Optical blur was also significantly correlated with response strength in both pathways (Figure 3e, R^2^=0.92, p=0.0035 for ON and R^2^=0.82, p=0.0259 for OFF), and temporal suppression in ON pathways (Figure 3f, R^2^=0.88, p=0.0099 for ON and R^2^=0.47, p=0.2904 for OFF).

**Figure 3.**
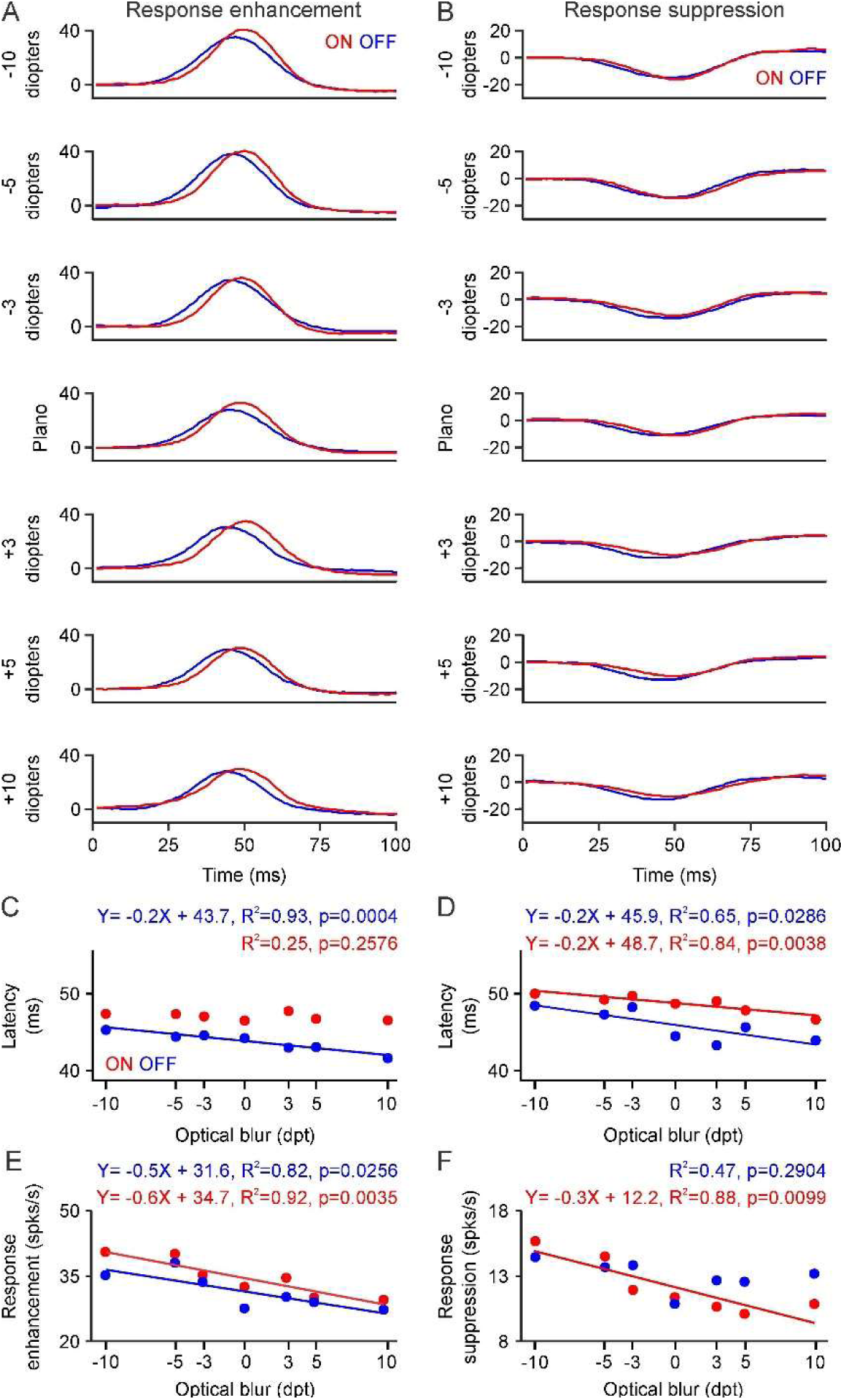
Positive spectacle lenses decrease cortical response latency. **(A)** Average response time-course to the ten preferred stimuli from a sequence of large grating patterns while inducing optical blur with spectacle lenses (ON-dominated neurons in red, n=327; OFF-dominated neurons in blue, n=284). **(B)** Same as A, but for response suppression to the ten non-preferred gratings. **(C)** Average peak response latency to the preferred gratings for ON-and OFF-dominated neurons while inducting optical blur with spectacles (top labels: R^2^, p value of linear regression, and equation reported only for significant correlations). **(D)** Same as C for response suppression to non-preferred gratings. **(E-F)** Same as C-D for response amplitude. * p<0.05, ** p<0.01, *** p<0.001 (Wilcoxon tests). Error bars indicate ± SEM.

### ON and OFF human pathways have different spatial resolution

ON and OFF cortical pathways also differed in their preference for stimulus size. In cat visual cortex, ON pathways respond to larger stimuli than OFF pathways ^19,20^, replicating differences already present in the retina of a large variety of mammals including rodents, carnivores, non-human primates and humans ^12,36–38^. We replicate these ON-OFF differences in human visual cortex by measuring responses to the onset of light and dark checkerboard stimuli with electroencephalography. In our human cortical measures, small stimuli drove stronger responses from OFF than ON pathways while large surfaces drove stronger responses from ON than OFF pathways (Figure 4). These ON-OFF differences could be demonstrated in both individual subjects (Figure 4a-b) and the subject average (Figure 4c). On average, responses to small 0.14° stimuli were 23% stronger in OFF than ON pathways (Figure 4c, OFF: 0.921±0.029, ON: 0.746±0.052, n=18, p=0.0002, Wilcoxon test), whereas responses to large surfaces were 17-43% stronger in ON than OFF pathways. The stronger ON pathway responses to large surfaces could be demonstrated with check stimuli that were three, four and five times larger than the optimal stimulus size (OFF/ON: 0.687±0.060/0.801±0.054 for 0.58°, p=0.0299; 0.527±0.052/0.679±0.054 for 1.12°, p=0.0033;0.408±0.038/0.583±0.051 for 2.24°, p=0.0002, n=18 subjects, Wilcoxon tests).

**Figure 4.**
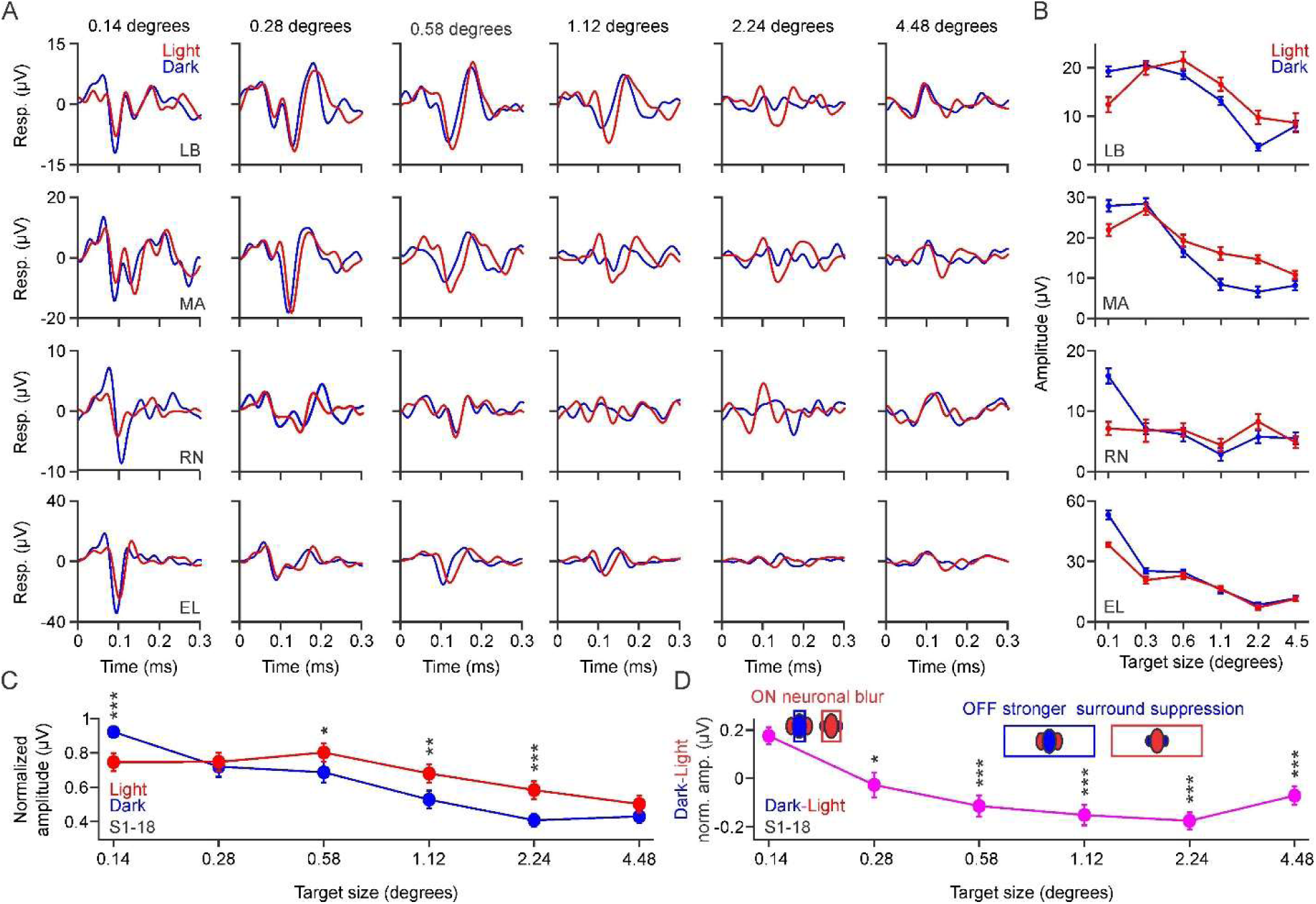
Human cortical responses show a preference for small dark stimuli and large light surfaces. **(A)** Human cortical responses to the onset of dark (blue) and light (red) checkerboards (0.5 sec, midgray background) measured with electroencephalography. Each row illustrates a different observer (LB, MA, RN, EL) and each column a different check size (from 0.14 to 4.48 degrees). **(B)** Size tuning for dark and light checkerboard stimuli calculated as the maximum minus minimum response between 50 and 200 ms after response onset (25-75 repeats). **(C)** Average (n=18) normalized cortical responses to dark (blue) and light stimuli (red) with different sizes (left). **(D)** Normalized dark minus light response. Top cartoons: small stimuli drive OFF pathways better (left) because light stimuli are more expanded by neuronal blur within the suppressive receptive field flanks. Large surfaces drive ON pathways better because OFF pathways have strong surround suppression.

Responses to stimuli smaller than the receptive field center are too weak to be measured with electroencephalography but should be stronger in ON than OFF cortical pathways because the contrast-response saturation of ON pathways (neuronal blur) expands light stimuli ^10,13,26,29,35^. Therefore, our results indicate that cortical responses are stronger in ON than OFF pathways when the stimulus is either very small (smaller than the receptive field center) or very large (larger than the receptive field surround). When the stimulus is smaller than the receptive field center, the greater contrast saturation of ON pathways (neuronal blur) expands the size of light more than dark stimuli making ON pathway responses stronger. When the stimulus size matches the receptive-field center, the greater contrast saturation of ON pathways expands the size of light stimuli within the suppressive surround making ON pathway responses weaker. When the stimulus size becomes several times larger than the entire receptive field (including center and surround), the responses are stronger in ON than OFF pathways because ON pathways have weaker surround suppression than OFF pathways ^11,19,20^ (Figure 4d).

### ON and OFF human cortical pathways are differently tuned to optical blur

We have previously demonstrated that optical blur decreases the visual salience of light more than dark stimuli in human vision ^8^. This light-dark difference can be explained by the expansion of ON receptive fields, which decreases spatial resolution for light more than dark stimuli. It can also be explained by the contrast loss associated with optical blur, which weakens surround suppression in ON more than OFF receptive fields ^19,20^. To our surprise, optical blur had the opposite effect on the strength of cortical responses. It weakened cortical responses to dark more than light stimuli (Fig 5, a-b). At focus, cortical responses were ∼20% stronger for dark than light stimuli (Figure 5c, top, darks/Lights: 0.797±0.063%, n=10, p=0.0313, Wilcoxon test), as expected from the cortical dominance of OFF pathways ^13–15,17,18,24,29,30,32–34^. However, optical blur reduced this OFF dominance. On average, cortical responses measured with electroencephalography under optical blur were ∼120 % stronger to light than dark stimuli (Figure 5c, top, dark vs. light: 0.414±0.050 vs. 0.829±0.069 at -5 diopters, p=0.002; 0.342±0.042 vs. 0.813±0.063 at +5 diopters, p=0.002, n=10 subjects, Wilcoxon tests), and the OFF-ON response difference became more negative as the optical blur increased (Figure 5c, bottom). The effect of optical blur was also more diverse for light than dark stimuli. Across subjects, optical blur consistently decreased cortical responses to dark stimuli, but could make the responses to light stimuli stronger (Figure 5a-b, LB and RN), relatively unchanged (Figure 5a-b, EL and MA), or weaker (Figure 5a-b, AA and SP).

**Figure 5.**
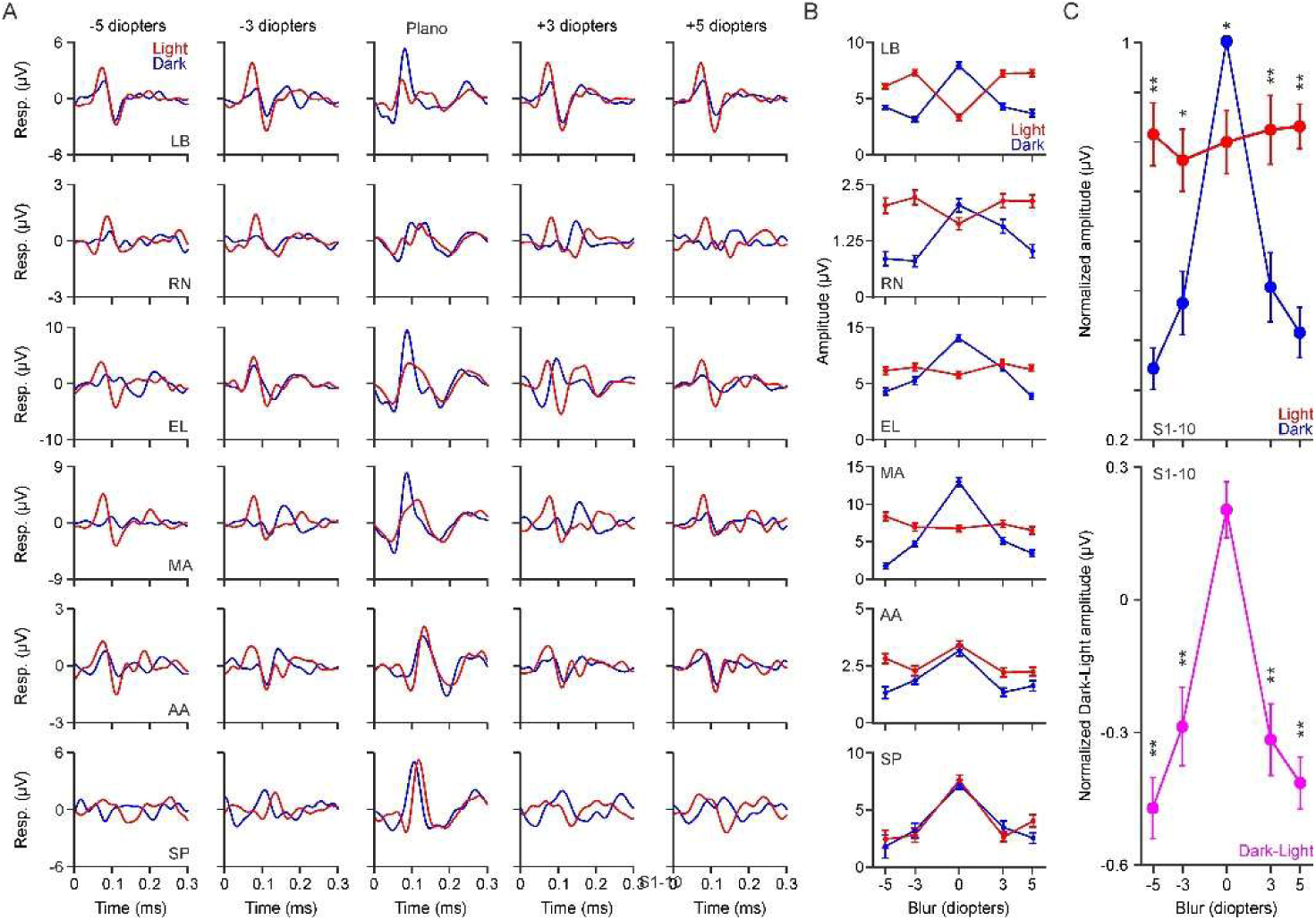
Blur tuning of ON and OFF pathways in human visual cortex. **(A)** Human cortical responses to the onset of dark (blue) and light (red) checkerboards measured with electroencephalography under different levels of optical blur induced with contact lenses. Each row illustrates an observer (LB, RN, EL, MA, AA, SP) and each column a level of optical blur (from -5 to +5 diopters). **(B)** Blur tuning for dark (blue) and light (red) stimuli calculated as the maximum minus minimum response between 50 and 200 ms after response onset. **(C)** Average (n=10) normalized cortical responses to dark and light stimuli measured with different levels of optical blur (top) and the normalized OFF minus ON response (magenta, bottom). Response amplitude is calculated as in B. ** p<0.01, *** p<0.001 (Wilcoxon tests). Error bars indicate ± SEM.

### Correlation between ON pathway blur tuning and response strength

The diversity of blur tuning measured with light stimuli could reflect individual differences in the population receptive field of ON pathways measured with electroencephalography. By shrinking dark stimuli, optical blur should consistently reduce the stimulus spatial summation and weaken OFF pathway responses. However, by expanding light stimuli, optical blur could make ON pathway responses stronger when the expansion is restricted to the receptive field center (Figure 6a, top), weaker when the expansion reaches the suppressive surround (Figure 6a, middle), or unchanged when the expansion is beyond the receptive field surround (Figure 6a, bottom). The strength of the ON pathway response at focus should be also different in these three conditions. It should be strongest when the stimulus is restricted to the receptive field center and weakest when it covers also the surround. Therefore, if our prediction is correct, ON pathway blur tuning should be correlated with ON pathway response strength.

**Figure 6.**
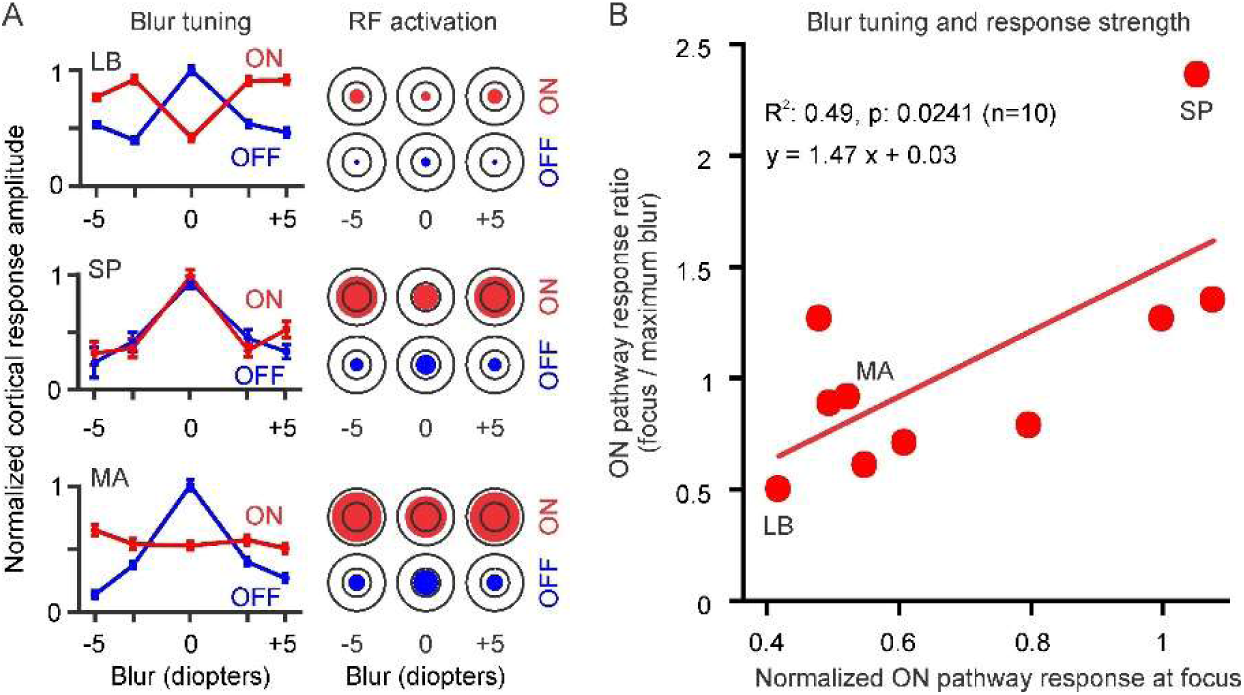
Correlation between ON pathway blur tuning and response strength. **(A)** Examples of three types of ON pathway blur tuning: U shape (LB), inverted U shape (SP), and flat (MA). The right of the figure panel illustrates how differences across subjects in population receptive field (RF) size could generate different blur tuning. The shrinkage of dark stimuli (blue) makes the activation of OFF receptive fields always weaker in the presence of optical blur because it reduces spatial summation at the receptive field center. However, the expansion of light stimuli (red) can make the activation of ON receptive fields more effective if the stimulus is smaller than the receptive field center (LB), less effective if the stimulus is roughly equal to the receptive field center (SP), and equally effective if the stimulus is larger than the receptive field center at focus (MA). The cortical population receptive field is concentric (see Mazade et al., 2022, Figure 2). **(B)** Correlation between ON pathway response strength (x axis) and blur tuning (y axis). Response strength is normalized by the OFF pathway response (ON/OFF ratio). The blur tuning is measured as the response at focus divided by the response at maximum blur. The circles show data from the ten observers of Figure 5 (LB, SP and MA also illustrated in panel A) and the line shows a linear regression (top: R^2^, p value, n, and equation).

To test this prediction, we measured ON pathway blur tuning by calculating a ratio between the response at focus and at maximum blur. We then measured ON pathway response strength at focus and normalized it by the OFF pathway response to eliminate variability caused by factors not relevant to the study (e.g. skull thickness, scalp impedance). Consistent with our prediction, our results demonstrate a significant correlation between ON pathway blur tuning and response strength (Figure 6b, R²=0.49, p=0.0241, n=10). As ON pathway responses became stronger at focus, optical blur was more effective at reducing response strength. We also found a significant correlation between ON pathway blur tuning and optimal stimulus size (R²=0.70, p=0.0389, n=6, range and median optimal size: 0.14° to 2.24° and 0.43°).

## DISCUSSION

Our results demonstrate that optical blur affects differently the receptive field properties and cortical responses of ON than OFF visual pathways. We show that the light scatter expands ON receptive fields while shrinking OFF receptive fields, making the population response weaker in OFF than ON pathways. We also demonstrate that the image magnification from positive spectacles decreases the receptive field size, response latency and response strength of OFF more than ON pathways. These results indicate that optical blur disrupts the response balance between ON and OFF pathways and the balance is not fully restored by correcting the refractive error with spectacles. Taken together with previous work, these results shed new light on the role of ON and OFF pathway function in signaling optical blur and refractive error ^8–10,39–42^.

### Differences in spatial resolution between ON and OFF pathways

In mammals with good visual acuity, OFF retino-thalamic pathways have smaller receptive fields, smaller dendritic fields, higher cell density, and higher receptive-field coverage than ON retino-thalamic pathways ^12,36–38^. These differences in spatial resolution are reliably transmitted to visual cortex, where they are amplified by the stronger surround suppression ^11,19,20^ and more precise retinotopic mapping of OFF than ON pathways ^29,30^. Our measurements in human visual cortex replicate these ON-OFF differences in spatial resolution previously demonstrated in carnivores and non-human primates. Consistently with the stronger responses and larger number of OFF than ON cortical receptive fields ^13,15–17,19,20,24,29–31,33,35,43^, small dark stimuli drove stronger visual responses than small light stimuli in human cortex (see also ^26,34,44^). Also, consistently with the stronger surround suppression of OFF pathways ^11,19,20^, large bright surfaces drove stronger responses than large dark surfaces.

### Processing of optical blur in ON and OFF pathways

Across the animal kingdom, visual spatial resolution is maximized by making eyes larger and retinal cell density higher ^45,46^. Larger eyes generate larger retinal images that are sampled more finely with a larger number of cells. Maximizing spatial resolution also requires a precise adjustment of eye size and optical power to accurately focus visual targets at different viewing distances. Animals with binocular foveal vision explore their visual environments with coordinated changes in optical power (accommodation) and eye vergence, which require an extensive network of neurons in the reticular formation, superior colliculus, cerebellum and cerebral cortex^47–50^.

Focusing images at the point of fixation also generates a gradient of optical blur across the retina ^51,52^ that could potentially be used to compute visual depth ^53,54^. Our results indicate that this blur gradient affects differently ON and OFF pathways. The light scatter expands ON receptive fields while shrinking OFF receptive fields, explaining the different effects of blurred vision on the perception of light and dark stimuli ^8,55^. The contrast reduction of optical blur also reduces the responses from OFF more than ON pathways at most spatial frequencies because ON pathway responses saturate more with contrast ^8,12,18,21,26,27^. Only the contrast loss at the highest spatial frequencies affects ON more than OFF pathways because ON pathways respond to higher spatial frequencies than OFF pathways ^13,18^.

### The importance of image magnification

Our results also indicate that, without changes in image size or luminance, the population response of ON and OFF pathways cannot reliably distinguish negative from positive blur. When induced with spectacles, optical blur is strongly correlated with OFF receptive field size but the correlation is caused by changes in image magnification, not blur polarity. When induced with contact lenses, negative blur expands ON receptive fields slightly more than positive blur. However, the difference is very small (0.28°, ∼5% of a receptive field for 20 diopters difference) and could be caused by an equally small difference in image magnification (∼6% calculated as 1 / (1-(p x d)), where p is -10 or +10 diopters and d is 3.2 mm of anterior chamber depth plus contact lens thickness).

ON and OFF pathways cannot discriminate blur polarity but can reliably signal changes in blur magnitude and image magnification. Increasing optical blur weakens the visual responses of OFF more than ON pathways because the light scatter shrinks OFF receptive fields while expanding ON receptive fields (Figure 7a-c). Moreover, because OFF pathways have stronger surround suppression than ON pathways ^11,19,20^, the image magnification affects OFF pathways more, making spectacle power strongly correlated with both OFF receptive field size (R^2^=0.98) and response strength (R^2^=0.82; responses to gratings are strongly OFF dominated ^13,19^). Image magnification also increases with eye axial length, which can vary in adult humans from 21.5 to 37 millimeters (∼53% range calculated as range/average ^56^). However, as the eye becomes longer, the background of the retinal image also becomes dimmer. For example, a ∼53% increase in eye axial length is associated with a ∼56% increase in image magnification (magnification = 0.01306 x (L-1.82) ^57^) and a ∼99% decrease in median luminance (light density = 1/L^2^, inverse square law; L is eye axial length). And these percentages are even larger if they include newborn eyes (axial length ∼16.5 mm ^58^). Because ON pathways respond stronger than OFF pathways to large stimuli, dark backgrounds, and low contrasts ^19,26^, increasing eye axial length should strengthen responses in ON more than OFF pathways.

**Figure 7.**
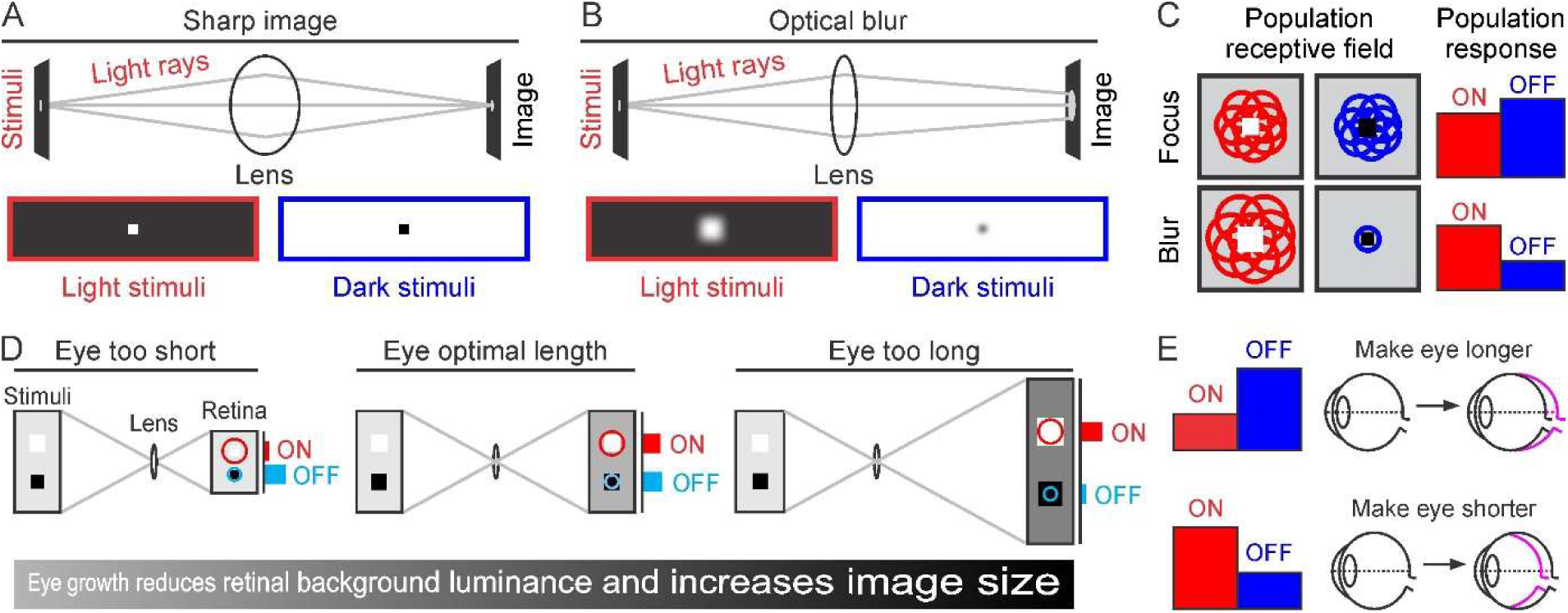
ON and OFF pathway function and retinal image quality. **(A)** A stimulus (light dot) generates divergent light rays (gray lines) that are focused by the crystalline lens onto the retina. Light stimuli on black background (bottom, red outline) appear slightly larger than dark stimuli on bright background (bottom, blue outline). **(B)** When the lens fails to converge the light rays, the light scatter expands light dots in black backgrounds (bottom, red outline) while shrinking dark dots in bright backgrounds (bottom, blue outline). **(C)** At focus (top), the population receptive field is just slightly smaller (left) and the population response just slightly stronger (right) in OFF than ON pathways. Optical blur (bottom) makes these ON-OFF differences opposite more pronounced (cartoon illustrates the size and number of receptive fields activated by the stimulus). **(D)** Light and dark stimuli on a midgray background drive different ON/OFF response balance when the eye is too short (left) or too long (right) than when it has optimal length (middle). As the eye grows (left to right), the retinal background luminance decreases and the image size increases. At the optimal eye length (middle), the combination of background luminance and image size is optimal to generate the maximum response from both ON and OFF pathways. **(E)** According to this mechanism, the eye increases in length (myopia) when the visual environment and/or genetic conditions do not drive effectively ON pathways (top). Conversely, the eye length should decrease (hyperopia) when ON pathways are overstimulated.

### An ON-OFF neuronal mechanism to optimize eye axial length

These results suggest that eye axial length could be accurately adjusted by optimizing the image brightness/size that drives the most balanced and strongest response from ON and OFF visual pathways. An eye that is too short should generate retinal images that are too small, have too bright background, and drive OFF better than ON pathways (Figure 7d, left) because OFF receptive fields are smaller and better activated under bright backgrounds than ON receptive fields ^8,12,13,18–20,37^. Conversely, an eye that is too long should generate retinal images that are too large, have too dark background, and drive ON better than OFF pathways (Figure 7d, right) because ON receptive fields are larger, have weaker surround suppression ^11,19,20^, and are better activated on dark backgrounds than OFF receptive fields (Figure 7d-e). Therefore, the eye should become longer when ON pathway responses are weak and shorter when ON pathway responses are strong (Figure 7e).

Because the responses from both ON and OFF pathways increase with luminance range ^10,19,26^, bright environments should drive both pathways very strongly (Figure 7e), and could explain why spending time outdoors protects against developing myopia ^59^ (notice that population cortical responses at focus are stronger in OFF than ON pathways ^17,24,34^). Conversely, because ON pathways are more vulnerable to reductions in luminance range ^10,18,26,60^, dim mesopic environments or frosted lenses should reduce responses in ON pathways and elongate the eye ^10,39,40,42^. Negative blur ^61^ and reading ^39,42^ could also induce myopia through a reduction in luminance range caused by pupil constriction, which is part of the accommodation reflex compensating negative blur (unlike pupil constriction, eye elongation preserves luminance range). Conversely, atropine would protect against myopia progression by increasing the retinal illumination with pupil dilation ^62^ and, in animals that lack accommodation and/or have small pupils (e.g. mice), inducing myopia should require powerful negative lenses with pronounced image minification. Because all animals with form vision have ON and OFF pathways ^45,63^, this ON-OFF mechanism could be potentially implemented in a large variety of species. Therefore, by measuring the neuronal effects of optical blur, our work may have revealed a new role of ON and OFF pathways in maximizing the quality of the retinal image and guiding eye growth.

## Acknowledgements

Supported by NIH grant EY027361

## AUTHOR CONTRIBUTIONS

C.P., M.D., and J.-M.A. recruited the human observers. C.P., R.M., J.J. and J.-M.A. performed the electrophysiological measurements. C.P., J.J. and J.-M.A. performed the data analysis. The paper was written by C.P., R.M. and J.-M.A. and edited by all the authors.

## STAR* METHODS

### EXPERIMENTAL MODEL AND STUDY PARTICIPANT DETAILS

#### Electrophysiological recordings in cat visual cortex

Adult male cats (Felis catus, 4-7 kg, n=12) were housed in groups of 2-3 animals in the biological research facility where they had free roaming time, and daily interaction with humans. They were also provided with enrichment toys and Purina cat food in their private room. All procedures performed on animals were approved by the Institutional Animal Care and Use Committee (IACUC) at the State University of New York, College of Optometry, and were performed in accordance with the guidelines of the United States Department of Agriculture (USDA). The animals were first tranquilized with an intramuscular injection of acepromazine (0.2 mg/kg) and ketamine (10 mg/kg). Following the intramuscular injection, two intravenous catheters were placed into each hind limb for continuous infusions of propofol (3–6 mg/kg/h), sufentanil (10–20 ng/kg/1), vecuronium bromide (0.2 mg/kg/h) and normal saline (1–3 ml/h). During the experiment, heart rate, blood pressure, electrocardiogram, temperature, pulse oximetry, expired CO2 and electroencephalographic activity were carefully monitored to keep the values within normal physiological limits. The nictitating membranes were retracted with 2% neosynephrine and the pupils dilated with 1% atropine sulfate. The eyes were fitted with contact lenses that had an artificial pupil of 3 mm to protect the corneas and focus the stimuli on the retina. In some experimental conditions, contact lenses with different optical power were also used to induce optical blur on the retinal image. Similar surgical procedures have been previously described in further detail ^19,26^.

#### Human electroencephalographic (EEG) recordings

We performed EEG recordings in 23 human observers (12 females and 11 males, age range: 20-55 years old, including three authors C.P., R.M. and J.M.A). 21 observers participated in the measurements of size tuning (12 females and 9 males, age range: 20-55 years old), 8 of these 21 observers participated also in the measurements of blur tuning (6 females and 2 males, age range: 20-29 years old), and 2 observers participated only in the measurements of blur tuning (2 males, age range 25-27 years old). 3 observers that participated in the measurements of size tuning were not selected for further analysis because their maximum cortical responses were very weak (< 12 microvolts, 1 female and 2 males including one author: J.M.A.). Therefore, size tuning was measured in 18 observers (11 females and 7 males, age range: 20-50 years old, age mean: 28.3 years old), and blur tuning was measured in 10 observers (6 females and 4 males, age range: 20-43 years old, age mean: 27.1 years old). All recruited observers had 20/20 vision (or vision at focus corrected to 20/20). All observers that participated on the blur tuning experiments had a non-corrected visual acuity between 20/15 and 20/30 and no differences in visual acuity between the eyes. All experiments in human subjects were approved by the institutional review board at the State University of New York College of Optometry and followed the principles outlined in the Declaration of Helsinki.

### METHOD DETAILS

#### Cortical recordings in cat visual cortex

Multiunit cortical activity was measured with 32-channel linear multielectrode arrays (0.1 mm inter-electrode distance, Neuronexus) with one, two or four shanks (200-400 microns of inter-shank distance). The multielectrodes were introduced in cat primary visual cortex nearly parallel to the cortical surface and centered in layer 4 (around 2 mm lateral to the brain midline and 4-5 mm posterior to stereotaxic zero). At the cortical locations recorded, most receptive fields were within 10 degrees of eccentricity, between the horizontal meridian and the lower visual field. The electrophysiological recordings were collected with a computer running Omniplex (Plexon), filtered between 250 Hz and 8 kHz and sampled at 40 kHz as previously described ^17,19,26^. Custom code with MATLAB (Mathworks) and the Psychtoolbox extension ^64^ was used to present visual stimuli on two gamma corrected monitors, a 24 inch LCD monitor (BenQ XL2420-B, 120 Hz, mean luminance: 120 cd/m²) and a 24 inch VIEWPixx monitor (VIEWPixx /3D, 120 Hz, mean luminance: 48.2 cd/m²). The monitors were placed at a distance of 0.57 meters from the eye. The receptive field measurements with sparse noise stimuli were all obtained with the LCD monitor. The measurements with a fast sequence of gratings were obtained with both LCD and VIEWPixx monitors.

Optical blur was induced with both contact lenses and spectacles. To induce optical blur with contact lenses, we fitted contact lenses of -10 or +10 diopters to the eye. To induce optical blur with spectacles, the animals were fit with contact lenses of the adequate diopter power to focus the image on the retina and then spectacles of different power were placed in front of the eye at ∼ 1 cm distance to induce optical blur with -10, -5, -3, +3, +5, or +10 diopters. We measured receptive field diameter, firing rate and peak response latency for each level of optical blur.

#### Data acquisition of human cortical responses

Human cortical responses were measured with a custom dry electrode wireless electroencephalogram (EEG) system (Wearable Sensing DSI-7-Flex, San Diego, CA) that allowed us to sample the human visual cortex with the same density than a 128-channel EEG. The device had 7 electrodes centered on the occipital cortex of our human observers according to the “10-20 International System” (Oz, O1, O2, Poz, Po3, Po4, and Fpz, impedances < 1 MΩ). The Oz electrode was positioned 3–4 cm above the inion and the ground, consisting of 3 separate electrodes, was positioned across the forehead. EEG signals were sampled at 600 Hz, low-pass filtered at 100 Hz, and collected by a computer running Rasputin (Plexon).

Human observers were asked to fixate on a small green dot centered on a computer monitor with one eye while the contralateral eye was covered by a patch. After fixation, observers were instructed to refrain from blinking for 6.6 seconds of stimulus presentation. Eye movements and pupil size were continuously monitored with an EyeLink 1000 Plus (SR Research Ltd, Ottawa, ON) and the trials were aborted and repeated if the observer blinked or ceased fixation in the middle of the trial. To induce optical blur, accommodation was blocked with cyclopentolate (1%) and contact lenses were placed on the test eye of the observer. The contact lenses were tested in the following order: 0, -3, -5, +3 and +5 diopters. For each diopter level and stimulus size, dark stimuli were tested before light stimuli. The pupil size remained roughly constant across stimulus conditions and blur levels as expected from the constant background (measurements of size tuning) or the inactivation of the pupillary reflex with 1% cyclopentolate (measurements of blur tuning).

#### Presentation of visual stimuli for human subjects

We generated custom code with MATLAB (Mathworks) and the Psychtoolbox extension (Brainard, 1997) to present visual stimuli on a gamma corrected 24-inch VIEWPixx monitor (VIEWPixx /3D, 120 Hz, mean luminance: 48.2 cd/m²) that was placed at a distance of 1 meter from the observer’s eye. To measure human EEG responses, we presented checkerboards that consisted of light (96.48 cd/m²), dark (0.46 cd/m²) and gray (48.2 cd/m²) squares and that were turned on for 0.5 seconds and turned off for 0.6 seconds on each stimulus cycle. Because the signal to noise ratio is much lower in electroencephalographic recordings than in recordings of cortical multiunit activity, the human cortical responses were measured with checkerboard stimuli instead of white noise or static gratings. To measure the effect of stimulus size on electroencephalographic responses to light and dark stimuli, we presented checkerboard patterns that ranged from 0.14 to 4.48 degrees per side. The stimulus presentation always followed the order: 4.48, 2.24, 1.12, 0.58, 0.28, 0.14 degrees. For each stimulus size, we tested dark stimuli before light stimuli. When the checkerboard was turned off, a uniform gray (48.2 cd/m²) background surface was shown. We presented 25-75 stimulus sequences, each containing 6 stimulus sizes. Each observer performed 300-900 trials (25-75 trials x 2 polarities x 6 sizes) that took about 15-30 minutes of visual stimulation. A 5 -10 min break was given every 150 trials.

To measure the effect of optical blur, we presented checkerboard patterns of 0.28 degrees per side. When the checkerboard was turned off, a uniform light (96.48 cd/m²) or dark (0.46 cd/m²) background surface was shown. We presented 25 stimulus sequences, each containing 3 stimulus cycles per luminance range. The checkerboards were presented in a sequence of 75 trials per luminance range. We measured cortical responses to light (96.48 cd/m²), dark (0.46 cd/m²) and gray (48.2 cd/m²) stimuli under five different levels of optical blur (-5, -3, 0, 3, and 5 diopters). Each observer performed 1500 trials (75 trials x 2 polarities x 2 luminance levels x 5 optical blur levels) that took about 75 minutes of visual stimulation. A 5-10 min break was given every 150 trials.

#### Code accessibility

Custom code requests can be directed to the Lead Contact, Jose-Manuel Alonso.

### QUANTIFICATION AND STATISTICAL ANALYSIS

#### Analysis of receptive fields in cat visual cortex

The receptive fields were mapped by spike-trigger averaging sparse light (239 cd/m²) and dark (0.27 cd/m²) square stimuli presented on a light (239 cd/m²) or dark (0.27 cd/m²) background with an update rate of 30 Hz in the LCD monitor (20 × 20 target positions, separation between positions: 1.4 degrees, target width: 2.8 degrees). The receptive field diameter and mean firing rate were measured at the response peak with a threshold of 20% of the maximum response. We also calculated the peri-stimulus time histograms (PSTHs) from visual responses to stationary gratings of 88 different orientations, 41 different spatial frequencies and 4 different phases presented at 60 Hz ^65^. The responses to these stationary gratings were also used to quantify the average peak latency to preferred and non-preferred stimuli.

We classified the selected receptive fields based on their ON/OFF dominance, defined as the contrast polarity at the time interval in which the receptive field reached its largest absolute value. Contrast polarity was calculated as (ON-OFF)/(ON+OFF), where ON and OFF are the maximum ON and OFF responses measured at the selected time frame. After baseline-subtracting and normalizing the receptive field, we calculated the receptive field diameter in degrees as sqrt (2 * pixel count / pi). We also measured visual responses to sparse noise with PSTHs calculated with 1 ms bins for light (239 cd/m²) or dark (0.27 cd/m²) squares located at the center of each receptive field. The sparse-noise PSTHs were baseline subtracted and smoothed with a temporal window of 5 ms. The signal-to-noise ratio of sparse-noise PSTHs was measured separately for each recording site and level of optical blur. Blur-tuning measurements were obtained for -10, -5, 0, +5, and +10 diopters for spectacles and -10, 0 and +10 diopters for contact lenses.

Blur-tuning was measured in sparse-noise PSTHs that fulfilled the following criteria: 1) The PSTH averaged across all blur levels had a signal-to noise ratio larger than 5 when blur was induced with contact lenses and larger than 10 when induced with spectacles. 2) The weakest PSTH response generated by a given blur level had a signal-to-noise ratio larger than 3 when blur was induced with contact lenses and larger than 5 when induced with spectacles. 3) The onset responses were larger than the rebound responses for all blur levels (onset response: response to stimulus onset; rebound response: response to stimulus being turned off). 4) All blur levels generated a response larger than 0.1 spikes per second. 5) The R square of the Gaussian function fitted to the receptive field was higher than 0.75 for all blur levels. After selecting the measurements that passed these criteria, we averaged the receptive field diameters separately for each blur level. The signal to noise ratio of the sparse-noise PSTH was calculated as the maximum response divided by the baseline response. The maximum response was measured by averaging responses within 4 ms around the maximum. The baseline response was calculated by averaging responses between 0 and 40 ms following the stimulus onset.

#### Analysis of peak response latency in cat visual cortex

To measure the peak response latency, we presented static gratings on both the LCD (mean luminance: 120 cd/m²) and the VIEWPixx (mean luminance: 48.2 cd/m²) monitors. We used grating sequences made of 576 distinct light (239 cd/m² on the LCD or 96.48 cd/m² on the VIEWPixx) or dark (0.27 or 0.46 cd/m²) gratings of 24 degrees in size presented on a midgray background (120 or 48.2 cd/m²). The gratings had 88 different orientations, 41 different spatial frequencies, 4 different phases and were presented at 60 Hz. For each recording site, we selected the 10 gratings that drove the strongest response and plotted the average grating PSTH calculated between -50 and +400 ms from the stimulus onset ^66^. These grating PSTHs were calculated with 1 ms bins and smoothed with a temporal window of 20 ms using the MATLAB function ‘smooth’.

The PSTH average for the 10 preferred gratings was used to measure the response to the preferred stimulus phase in each multiunit recording. We also averaged the PSTHs for the 10 gratings with opposite phase to the 10 preferred gratings to measure the response to the non-preferred stimulus phase. For this analysis, we also subtracted the baseline and calculated a signal-to-noise ratio of the average PSTH separately for each blur level. The signal to noise ratio of the grating PSTH was calculated as the maximum response divided by the baseline. The baseline activity was calculated as the average spike rate between -40 and 0 ms of the stimulus onset. For each blur level, the measurements of peak latency for the preferred grating phase and valley latency for the non-preferred grating phase were calculated only for PSTHs with a signal-to-noise ratio larger than 2.5.

#### Analysis of firing rate in cat visual cortex

Firing rate was calculated in two different ways. We first measured the mean firing rate in response to light (239 cd/m²) or dark (0.27 cd/m²) squares presented by the LCD monitor. We also measured firing rate from visual responses to a fast sequence of static grating stimuli presented by both the LCD and the VIEWPixx monitors. Mean firing rate for each recording site was calculated as the number of spikes during the time of stimuli presentation, which was approximately 5 minutes for both sparse noise and static gratings. Stimuli were presented under different levels of optical blur induced with contact lenses (-10, 0, +10 diopters) or with spectacles (-10, -5, -3, 0, +3, +5, +10 diopters). We calculated the mean firing rate for each blur level averaging firing rate of the same recording sites that passed our SNR criteria on the receptive field diameter and latency analysis.

#### EEG analysis in human subjects

To quantify the electroencephalographic activity, we first filtered the signal with a low-pass filter of 5 Hz and high-pass filter of 30 Hz cut-off frequency. For analysis of size tuning, we isolated the EEG signal coming from the central electrode, Oz. The single Oz signal gives the best representation of the size/response tuning in primary visual cortex, but the size tuning is greatly affected by the proximity of the EEG electrode to the cortical representation of the fovea, which varies across subjects. Of the 21 observers initially recruited, we selected the 18 whose maximum response amplitude was bigger than 12 microvolts. We then averaged the recordings from Oz over the 25-75 trials performed by the selected observers for each condition and calculated response amplitude as the difference between maximum and minimum response within 50 to 200 ms from the stimulus onset, where the signal to noise ratio of our stimuli was the highest. Response trials with an amplitude larger than 100 microvolts were classified as recording artifacts and not included in the analysis. For analysis of blur tuning, we performed a voltage subtraction across our electrode array to calculate the current density (D) between the 2 central electrodes, Oz and Poz. The voltage subtraction consisted of averaging the voltage of the peripheral electrodes (VO1, VPo1, VO2 and VPo2) and subtracting it from the average of the central electrodes (VOz+VPoz) as indicated on equation 1.

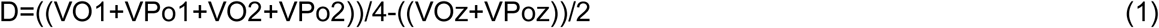

We then averaged the results of the voltage subtraction over the 75 trials performed by all the recruited observers for each condition and calculated response amplitude as the difference between maximum and minimum response within 50 to 200 ms from the stimulus onset, where the signal to noise ratio of our stimuli was the highest. We classified them as artifacts and did not include them in our analysis trials with an amplitude of more than 100 microvolts.

#### Statistical analyses for all experiments

We used the Wilcoxon Signed Rank tests for paired samples (MATLAB function ’signrank’) and Wilcoxon Ranked Sum tests for unpaired samples (MATLAB function ‘ranksum’) to calculate the significance of the differences between receptive field diameter, firing rate, peak latency and amplitude of the EEG signal within light and dark responses and within the different levels of optical blur. Error bars are ± standard error of the mean (SEM). Significance is noted as *p<0.05, **p<0.01, ***p<0.001.

## KEY RESOURCES TABLE

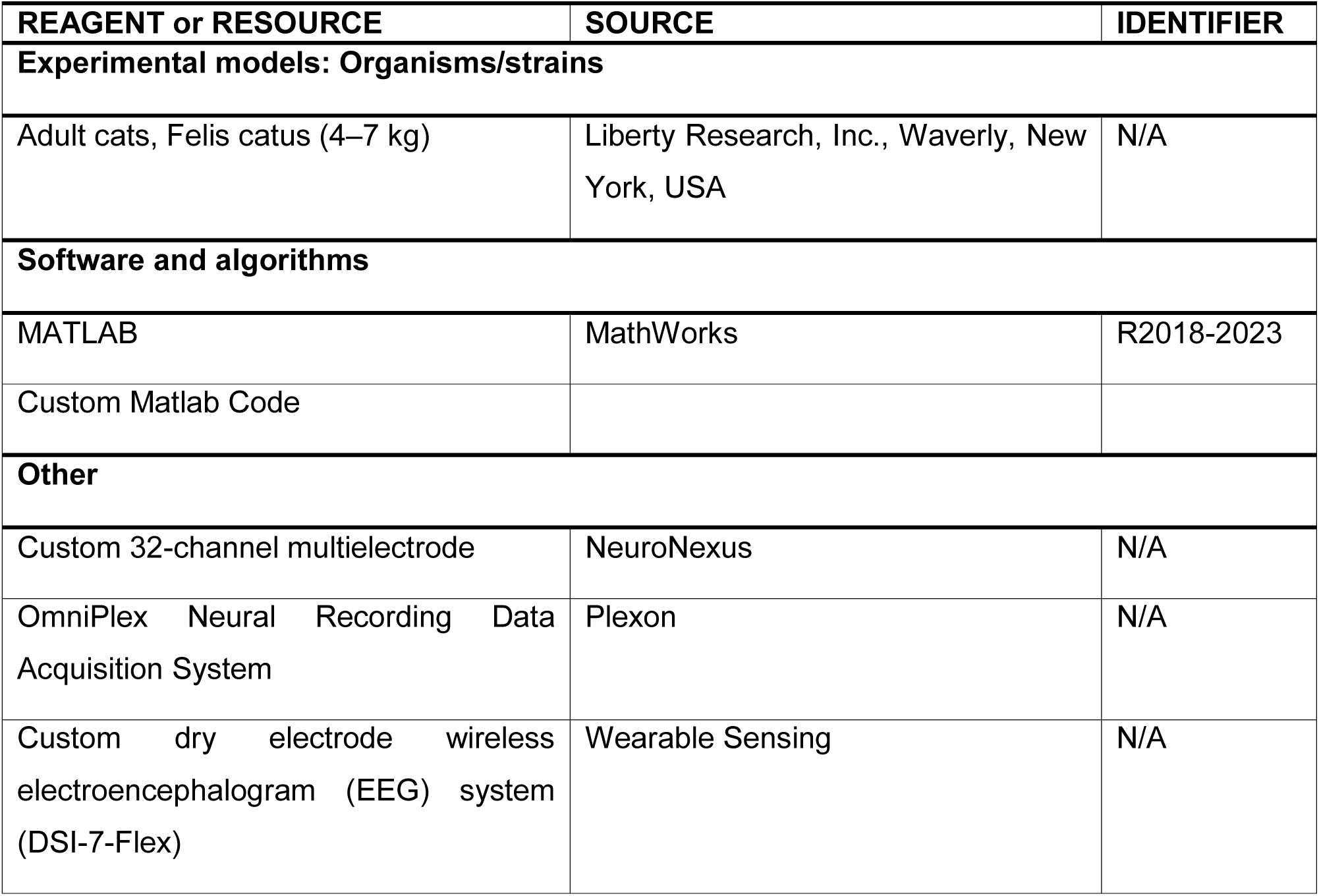

| REAGENT or RESOURCE | SOURCE | IDENTIFIER |
| --- | --- | --- |
| **Experimental models: Organisms/strains** |  |  |
| Adult cats, Felis catus (4–7 kg) | Liberty Research, Inc., Waverly, New York, USA | N/A |
| **Software and algorithms** |  |  |
| MATLAB | MathWorks | R2018-2023 |
| Custom Matlab Code |  |  |
| **Other** |  |  |
| Custom 32-channel multielectrode | NeuroNexus | N/A |
| OmniPlex Neural Recording Data Acquisition System | Plexon | N/A |
| Custom dry electrode wireless electroencephalogram (EEG) system (DSI-7-Flex) | Wearable Sensing | N/A |

## REFERENCES

1. Schaeffel, F., and Feldkaemper, M. (2015). Animal models in myopia research. Clin Exp Optom 98, 507–517. 10.1111/cxo.12312.

2. Wallman, J., Turkel, J., and Trachtman, J. (1978). Extreme myopia produced by modest change in early visual experience. Science 201, 1249–1251. 10.1126/science.694514.

3. Clavagnier, S., Dumoulin, S.O., and Hess, R.F. (2015). Is the Cortical Deficit in Amblyopia Due to Reduced Cortical Magnification, Loss of Neural Resolution, or Neural Disorganization? J Neurosci 35, 14740–14755. 10.1523/JNEUROSCI.1101-15.2015.

4. Hubel, D.H., Wiesel, T.N., and LeVay, S. (1977). Plasticity of ocular dominance columns in monkey striate cortex. Philos Trans R Soc Lond B Biol Sci 278, 377–409. 10.1098/rstb.1977.0050.

5. Kiorpes, L. (2006). Visual processing in amblyopia: animal studies. Strabismus 14, 3–10. 10.1080/09273970500536193.

6. Levi, D.M. (2006). Visual processing in amblyopia: human studies. Strabismus 14, 11–19. 10.1080/09273970500536243.

7. Wiesel, T.N., and Hubel, D.H. (1963). Single-Cell Responses in Striate Cortex of Kittens Deprived of Vision in One Eye. J Neurophysiol 26, 1003–1017. 10.1152/jn.1963.26.6.1003.

8. Pons, C., Mazade, R., Jin, J., Dul, M.W., Zaidi, Q., and Alonso, J.M. (2017). Neuronal mechanisms underlying differences in spatial resolution between darks and lights in human vision. J Vis 17, 5. 10.1167/17.14.5.

9. Pons, C., Jin, J., Mazade, R., Dul, M., Zaidi, Q., and Alonso, J.M. (2019). Amblyopia Affects the ON Visual Pathway More than the OFF. J Neurosci 39, 6276–6290. 10.1523/JNEUROSCI.3215-18.2019.

10. Poudel, S., Jin, J., Rahimi-Nasrabadi, H., Dellostritto, S., Dul, M.W., Viswanathan, S., and Alonso, J.M. (2024). Contrast Sensitivity of ON and OFF Human Retinal Pathways in Myopia. J Neurosci 44. 10.1523/JNEUROSCI.1487-23.2023.

11. Archer, D.R., Alitto, H.J., and Usrey, W.M. (2021). Stimulus Contrast Affects Spatial Integration in the Lateral Geniculate Nucleus of Macaque Monkeys. J Neurosci 41, 6246–6256. 10.1523/JNEUROSCI.2946-20.2021.

12. Chichilnisky, E.J., and Kalmar, R.S. (2002). Functional asymmetries in ON and OFF ganglion cells of primate retina. J Neurosci 22, 2737–2747. 10.1523/JNEUROSCI.22-07-02737.2002.

13. Jansen, M., Jin, J., Li, X., Lashgari, R., Kremkow, J., Bereshpolova, Y., Swadlow, H.A., Zaidi, Q., and Alonso, J.M. (2019). Cortical Balance Between ON and OFF Visual Responses Is Modulated by the Spatial Properties of the Visual Stimulus. Cereb Cortex 29, 336–355. 10.1093/cercor/bhy221.

14. Jimenez, L.O., Tring, E., Trachtenberg, J.T., and Ringach, D.L. (2018). Local tuning biases in mouse primary visual cortex. J Neurophysiol 120, 274–280. 10.1152/jn.00150.2018.

15. Jin, J., Wang, Y., Swadlow, H.A., and Alonso, J.M. (2011). Population receptive fields of ON and OFF thalamic inputs to an orientation column in visual cortex. Nat Neurosci 14, 232–238. 10.1038/nn.2729.

16. Jin, J., Wang, Y., Lashgari, R., Swadlow, H.A., and Alonso, J.M. (2011). Faster thalamocortical processing for dark than light visual targets. J Neurosci 31, 17471–17479. 10.1523/JNEUROSCI.2456-11.2011.

17. Jin, J.Z., Weng, C., Yeh, C.I., Gordon, J.A., Ruthazer, E.S., Stryker, M.P., Swadlow, H.A., and Alonso, J.M. (2008). On and off domains of geniculate afferents in cat primary visual cortex. Nat Neurosci 11, 88–94. 10.1038/nn2029.

18. Kremkow, J., Jin, J., Komban, S.J., Wang, Y., Lashgari, R., Li, X., Jansen, M., Zaidi, Q., and Alonso, J.M. (2014). Neuronal nonlinearity explains greater visual spatial resolution for darks than lights. Proc Natl Acad Sci U S A 111, 3170–3175. 10.1073/pnas.1310442111.

19. Mazade, R., Jin, J., Rahimi-Nasrabadi, H., Najafian, S., Pons, C., and Alonso, J.M. (2022). Cortical mechanisms of visual brightness. Cell Rep 40, 111438. 10.1016/j.celrep.2022.111438.

20. Mazade, R., Jin, J., Pons, C., and Alonso, J.M. (2019). Functional Specialization of ON and OFF Cortical Pathways for Global-Slow and Local-Fast Vision. Cell Rep 27, 2881–2894 e2885. 10.1016/j.celrep.2019.05.007.

21. Ravi, S., Ahn, D., Greschner, M., Chichilnisky, E.J., and Field, G.D. (2018). Pathway-Specific Asymmetries between ON and OFF Visual Signals. J Neurosci 38, 9728–9740. 10.1523/JNEUROSCI.2008-18.2018.

22. Rekauzke, S., Nortmann, N., Staadt, R., Hock, H.S., Schoner, G., and Jancke, D. (2016). Temporal Asymmetry in Dark-Bright Processing Initiates Propagating Activity across Primary Visual Cortex. J Neurosci 36, 1902–1913. 10.1523/JNEUROSCI.3235-15.2016.

23. Xing, D., Yeh, C.I., Gordon, J., and Shapley, R.M. (2014). Cortical brightness adaptation when darkness and brightness produce different dynamical states in the visual cortex. Proc Natl Acad Sci U S A 111, 1210–1215. 10.1073/pnas.1314690111.

24. Yeh, C.I., Xing, D., and Shapley, R.M. (2009). "Black" responses dominate macaque primary visual cortex v1. J Neurosci 29, 11753–11760. 10.1523/JNEUROSCI.1991-09.2009.

25. Zurawel, G., Ayzenshtat, I., Zweig, S., Shapley, R., and Slovin, H. (2014). A contrast and surface code explains complex responses to black and white stimuli in V1. J Neurosci 34, 14388–14402. 10.1523/JNEUROSCI.0848-14.2014.

26. Rahimi-Nasrabadi, H., Jin, J., Mazade, R., Pons, C., Najafian, S., and Alonso, J.M. (2021). Image luminance changes contrast sensitivity in visual cortex. Cell Rep 34, 108692. 10.1016/j.celrep.2021.108692.

27. Zaghloul, K.A., Boahen, K., and Demb, J.B. (2003). Different circuits for ON and OFF retinal ganglion cells cause different contrast sensitivities. J Neurosci 23, 2645–2654. 10.1523/JNEUROSCI.23-07-02645.2003.

28. Onat, S., Nortmann, N., Rekauzke, S., Konig, P., and Jancke, D. (2011). Independent encoding of grating motion across stationary feature maps in primary visual cortex visualized with voltage-sensitive dye imaging. Neuroimage 55, 1763–1770. 10.1016/j.neuroimage.2011.01.004.

29. Kremkow, J., Jin, J., Wang, Y., and Alonso, J.M. (2016). Principles underlying sensory map topography in primary visual cortex. Nature 533, 52–57. 10.1038/nature17936.

30. Lee, K.S., Huang, X., and Fitzpatrick, D. (2016). Topology of ON and OFF inputs in visual cortex enables an invariant columnar architecture. Nature 533, 90–94. 10.1038/nature17941.

31. Wang, Y., Jin, J., Kremkow, J., Lashgari, R., Komban, S.J., and Alonso, J.M. (2015). Columnar organization of spatial phase in visual cortex. Nat Neurosci 18, 97–103. 10.1038/nn.3878.

32. Williams, B., Del Rosario, J., Muzzu, T., Peelman, K., Coletta, S., Bichler, E.K., Speed, A., Meyer-Baese, L., Saleem, A.B., and Haider, B. (2021). Spatial modulation of dark versus bright stimulus responses in the mouse visual system. Curr Biol 31, 4172–4179 e4176. 10.1016/j.cub.2021.06.094.

33. Xing, D., Yeh, C.I., and Shapley, R.M. (2010). Generation of black-dominant responses in V1 cortex. J Neurosci 30, 13504–13512. 10.1523/JNEUROSCI.2473-10.2010.

34. Zemon, V., Gordon, J., and Welch, J. (1988). Asymmetries in ON and OFF visual pathways of humans revealed using contrast-evoked cortical potentials. Vis Neurosci 1, 145–150. 10.1017/s0952523800001085.

35. Komban, S.J., Kremkow, J., Jin, J., Wang, Y., Lashgari, R., Li, X., Zaidi, Q., and Alonso, J.M. (2014). Neuronal and perceptual differences in the temporal processing of darks and lights. Neuron 82, 224–234. 10.1016/j.neuron.2014.02.020.

36. Dacey, D.M., and Petersen, M.R. (1992). Dendritic field size and morphology of midget and parasol ganglion cells of the human retina. Proc Natl Acad Sci U S A 89, 9666–9670. 10.1073/pnas.89.20.9666.

37. Ratliff, C.P., Borghuis, B.G., Kao, Y.H., Sterling, P., and Balasubramanian, V. (2010). Retina is structured to process an excess of darkness in natural scenes. Proc Natl Acad Sci U S A 107, 17368–17373. 10.1073/pnas.1005846107.

38. Wassle, H., Boycott, B.B., and Illing, R.B. (1981). Morphology and mosaic of on-and off-beta cells in the cat retina and some functional considerations. Proc R Soc Lond B Biol Sci 212, 177–195. 10.1098/rspb.1981.0033.

39. Aleman, A.C., Wang, M., and Schaeffel, F. (2018). Reading and Myopia: Contrast Polarity Matters. Sci Rep 8, 10840. 10.1038/s41598-018-28904-x.

40. Chakraborty, R., Park, H.N., Hanif, A.M., Sidhu, C.S., Iuvone, P.M., and Pardue, M.T. (2015). ON pathway mutations increase susceptibility to form-deprivation myopia. Exp Eye Res 137, 79–83. 10.1016/j.exer.2015.06.009.

41. Chakraborty, R., Landis, E.G., Mazade, R., Yang, V., Strickland, R., Hattar, S., Stone, R.A., Iuvone, P.M., and Pardue, M.T. (2022). Melanopsin modulates refractive development and myopia. Exp Eye Res 214, 108866. 10.1016/j.exer.2021.108866.

42. Poudel, S., Rahimi-Nasrabadi, H., Jin, J., Najafian, S., and Alonso, J.M. (2023). Differences in visual stimulation between reading and walking and implications for myopia development. J Vis 23, 3. 10.1167/jov.23.4.3.

43. Taylor, M.M., Sedigh-Sarvestani, M., Vigeland, L., Palmer, L.A., and Contreras, D. (2018). Inhibition in Simple Cell Receptive Fields Is Broad and OFF-Subregion Biased. J Neurosci 38, 595–612. 10.1523/JNEUROSCI.2099-17.2017.

44. Norcia, A.M., Yakovleva, A., Hung, B., and Goldberg, J.L. (2020). Dynamics of Contrast Decrement and Increment Responses in Human Visual Cortex. Transl Vis Sci Technol 9, 6. 10.1167/tvst.9.10.6.

45. Mazade, R., and Alonso, J.M. (2017). Thalamocortical processing in vision. Vis Neurosci 34, E007. 10.1017/S0952523817000049.

46. Veilleux, C.C., and Kirk, E.C. (2014). Visual acuity in mammals: effects of eye size and ecology. Brain Behav Evol 83, 43–53. 10.1159/000357830.

47. Gamlin, P.D. (1999). Subcortical neural circuits for ocular accommodation and vergence in primates. Ophthalmic Physiol Opt 19, 81–89. 10.1046/j.1475-1313.1999.00434.x.

48. Gamlin, P.D., and Yoon, K. (2000). An area for vergence eye movement in primate frontal cortex. Nature 407, 1003–1007. 10.1038/35039506.

49. 49. Van Horn, M.R., Waitzman, D.M., and Cullen, K.E. (2013). Vergence neurons identified in the rostral superior colliculus code smooth eye movements in 3D space. J Neurosci 33, 7274–7284. 10.1523/JNEUROSCI.2268-12.2013.

50. Ward, M.K., Bolding, M.S., Schultz, K.P., and Gamlin, P.D. (2015). Mapping the macaque superior temporal sulcus: functional delineation of vergence and version eye-movement-related activity. J Neurosci 35, 7428–7442. 10.1523/JNEUROSCI.4203-14.2015.

51. Navarro, R., Artal, P., and Williams, D.R. (1993). Modulation transfer of the human eye as a function of retinal eccentricity. J Opt Soc Am A 10, 201–212. 10.1364/josaa.10.000201.

52. Wang, B., Ciuffreda, K.J., and Irish, T. (2006). Equiblur zones at the fovea and near retinal periphery. Vision Res 46, 3690–3698. 10.1016/j.visres.2006.04.005.

53. Banks, M.S., Sprague, W.W., Schmoll, J., Parnell, J.A., and Love, G.D. (2015). Why do animal eyes have pupils of different shapes? Sci Adv 1, e1500391. 10.1126/sciadv.1500391.

54. Mather, G., and Smith, D.R. (2002). Blur discrimination and its relation to blur-mediated depth perception. Perception 31, 1211–1219. 10.1068/p3254.

55. Sato, H., Motoyoshi, I., and Sato, T. (2016). On-Off asymmetry in the perception of blur. Vision Res 120, 5–10. 10.1016/j.visres.2015.03.010.

56. Panda-Jonas, S., Jonas, J.B., and Jonas, R.A. (2022). Photoreceptor density in relation to axial length and retinal location in human eyes. Sci Rep 12, 21371. 10.1038/s41598-022-25460-3.

57. Bennett, A.G., Rudnicka, A.R., and Edgar, D.F. (1994). Improvements on Littmann’s method of determining the size of retinal features by fundus photography. Graefes Arch Clin Exp Ophthalmol 232, 361–367. 10.1007/BF00175988.

58. 58. Groot, A.L.W., Lissenberg-Witte, B.I., van Rijn, L.J., and Hartong, D.T. (2022). Meta-analysis of ocular axial length in newborns and infants up to 3 years of age. Surv Ophthalmol 67, 342–352. 10.1016/j.survophthal.2021.05.010.

59. Rose, K.A., Morgan, I.G., Ip, J., Kifley, A., Huynh, S., Smith, W., and Mitchell, P. (2008). Outdoor activity reduces the prevalence of myopia in children. Ophthalmology 115, 1279–1285. 10.1016/j.ophtha.2007.12.019.

60. Barlow, H.B., and Levick, W.R. (1969). Changes in the maintained discharge with adaptation level in the cat retina. J Physiol 202, 699–718. 10.1113/jphysiol.1969.sp008836.

61. Schaeffel, F., Glasser, A., and Howland, H.C. (1988). Accommodation, refractive error and eye growth in chickens. Vision Res 28, 639–657. 10.1016/0042-6989(88)90113-7.

62. Chia, A., Chua, W.H., Cheung, Y.B., Wong, W.L., Lingham, A., Fong, A., and Tan, D. (2012). Atropine for the treatment of childhood myopia: safety and efficacy of 0.5%, 0.1%, and 0.01% doses (Atropine for the Treatment of Myopia 2). Ophthalmology 119, 347–354. 10.1016/j.ophtha.2011.07.031.

63. Kremkow, J., and Alonso, J.M. (2018). Thalamocortical Circuits and Functional Architecture. Annu Rev Vis Sci 4, 263–285. 10.1146/annurev-vision-091517-034122.

64. Brainard, D.H. (1997). The Psychophysics Toolbox. Spat Vis 10, 433–436.

65. Ringach, D.L., Sapiro, G., and Shapley, R. (1997). A subspace reverse-correlation technique for the study of visual neurons. Vision Res 37, 2455–2464. 10.1016/s0042-6989(96)00247-7.

66. Williams, P.E., and Shapley, R.M. (2007). A dynamic nonlinearity and spatial phase specificity in macaque V1 neurons. J Neurosci 27, 5706–5718. 10.1523/JNEUROSCI.4743-06.2007.

